# Interferon-induced activation state of circulating dendritic cells and monocytes triggered by yellow fever vaccination correlates with early protective antibody responses

**DOI:** 10.1101/2024.07.31.606034

**Authors:** Elena Winheim, Antonio Santos-Peral, Tamara Ehm, Linus Rinke, Sandra Riemer, Magdalena Zaucha, Sebastian Goresch, Lisa Lehmann, Katharina Eisenächer, Michael Pritsch, Giovanna Barba-Spaeth, Tobias Straub, Simon Rothenfusser, Anne B. Krug

## Abstract

Yellow fever vaccination provides long-lasting protection and is a unique model for studying the immune response to an acute RNA virus infection in humans. To elucidate the early innate immune events preceding the rapid generation of protective immunity, we performed transcriptome analysis of human blood dendritic cell (DC) and monocyte subpopulations before and 3, 7, 14 and 28 days after vaccination. We detected temporary upregulation of IFN-stimulated genes (ISG) in all DC and monocyte subsets on day 3 and 7 after vaccination as well as cell type specific responses and temporal dynamics. Single cell RNA sequencing revealed rapid appearance of activated DC and monocyte clusters dominated by ISGs, inflammatory chemokines and genes involved in antigen processing and presentation. This was confirmed by flow cytometric analysis in a large cohort of vaccinees. We identified SIGLEC1/CD169 upregulation as a sensitive indicator of the transient IFN-induced activation state elicited in DCs and monocytes by YF17D vaccination correlating with early protective IgM antibody responses.

## Introduction

Acute viral infections caused by flaviviruses such as yellow fever (YF), dengue and West Nile virus represent a major global health threat. The attenuated YF vaccine virus 17D-204 (YF17D) is the most successful live-attenuated vaccine available and provides long-lasting immune protection against yellow fever after a single dose (Fuertes Marraco et al., 2015; Gotuzzo et al., 2013; Kling et al., 2022; Poland et al., 1981). Therefore, this vaccine can be used as an excellent model to study the immunological characteristics of a highly efficient and long lasting protective immune response to an acute self-limited RNA virus infection.

The response to YF17D is marked by a systemic innate immune response involving the induction of and response to type I interferons, and inflammasome activation in peripheral blood mononuclear cells within the first week after vaccination (Gaucher et al., 2008; Hou et al., 2017; Querec et al., 2009). YF17D vaccination in rare incidents can cause life-threatening viscerotropic or neurotropic disease. In a recent study of 8 patients with severe vaccine-associated disease *IFNAR1* and *IFNAR2* deficiency as well as neutralizing autoantibodies against type I IFNs accounted for more than half of the cases, demonstrating the importance of the type I Interferon response in controlling YF17D replication and inducing protective immune responses (Bastard et al., 2021; Hernandez et al., 2019). Characteristically a transient activation of circulating dendritic cells (DCs) and monocytes within the first week after vaccination (Winheim et al., 2023) is followed by the activation and expansion of specific CD4^+^ and CD8^+^ T cells with a peak between 11 and 14 days after vaccination (Akondy et al., 2009; Huber et al., 2020; James et al., 2013). A strong humoral immune response with neutralizing antibodies initially dominated by IgM is detected within 2 weeks and persists in the majority of vaccinated patients (Lindsey et al., 2018; Poland et al., 1981; Santos-Peral et al., 2023). Data obtained in the mouse model suggest that the protective immune response to YF17D is mainly mediated by humoral immunity and CD4^+^ T cell responses (Watson et al., 2016), but cytotoxic CD8^+^ T cells are probably also relevant for long-term protection (Akondy et al., 2017). Systems vaccinology approaches using the transcriptome of whole PBMC defined genes and gene modules that may predict the adaptive immune response to YF17D and other vaccines. For example, expression of genes encoding molecules involved in the stress-response pathway such as *EIF2AK4* and B cell genes such as *TNFRSF17* were found to be predictive of the CD8^+^ T cell and antibody responses to YF17D respectively (Querec et al., 2009). Subsequently “blood transcriptional modules” were identified that correlated with antibody responses across several vaccines including YF17D (Hagan et al., 2022; Li et al., 2014). However, a comprehensive analysis of the responses of the different human DC and monocyte subpopulations to YF17D vaccination and their association with parameters of adaptive immunity is lacking until now.

Circulating monocytes rapidly respond to pathogens and pathogen-derived molecules by producing inflammatory mediators including cytokines and chemokines. Human monocytes comprise 3 subpopulations: CD14^+^ classical monocytes, CD14^+^ CD16^+^ intermediate monocytes and CD14^lo^ CD16^+^ non-classical patrolling monocytes. An increased frequency of proinflammatory CD14^+^ CD16^+^ intermediate monocytes has been observed in patients with bacterial sepsis, dengue fever, Zika virus and SARS-CoV2 infection (Fingerle et al., 1993; Kwissa et al., 2014; Michlmayr et al., 2017; Winheim et al., 2023) and also in response to YF vaccination (Martins et al., 2008; Winheim et al., 2023). DCs are professional antigen presenting cells (APCs) and essential for recognizing viruses and presenting viral antigens to T cells for induction of efficient adaptive immunity. In the human peripheral blood, we distinguish 3 types of conventional DCs (cDC1, cDC2, DC3), plasmacytoid DCs (pDC) and “transitional DCs” (tDCs) (See et al., 2017; Segura, 2022; Villani et al., 2017). While cDC1 preferentially promote cytotoxic T cell and Th1 responses, cDC2 are important for T helper (Th)2, Th17 and T follicular helper responses. DC3 which are phenotypically similar to cDC2 but also exhibit monocytic traits were shown to be proinflammatory and promote Th1 and Th17 responses and CD8^+^ tissue-resident memory T cells (Bourdely et al., 2020; Cytlak et al., 2020; Dutertre et al., 2019). CD14^+^ DC3 were described as an inflammatory DC subset that increases in frequency in the blood during flares of systemic lupus erythematosus (SLE) (Dutertre et al., 2019) and in COVID-19 patients correlating with disease severity (Winheim et al., 2021). pDCs produce large amounts of type I IFNs in response to viral stimulation. tDCs, also called pre-DCs or Axl+ Siglec 6+ (AS)-DCs (Alcántara-Hernández et al., 2017; See et al., 2017; Villani et al., 2017) can give rise to cDCs with cDC2 phenotype suggesting they are cDC2 precursors. At the same time, tDC actively participate in immune responses suggesting they have additional functions beyond their role as precursors (Sulczewski et al., 2023; Winheim et al., 2021). We hypothesized that functionally distinct DC and monocyte subpopulations show common as well as unique responses to YF17D vaccination and we sought to define cell-type specific activation states associated with effective and long-lasting adaptive immune responses in a vaccination cohort of naïve healthy adults.

## Results

### YF17D immunization induces a common core signature of IFN-stimulated genes in all APC subsets and cell-type specific responses

To characterize the response of circulating APC subpopulations to YF17D vaccination, bulk RNA sequencing was performed on sorted DC and monocyte subsets, B cells as well as total PBMCs, before and on days 3, 7, 14 and 28 after vaccination in four donors, 2 males and 2 females between 23 and 27 years of age (Figure 1A). Principal component analysis (PCA) of the cell subset transcriptomes showed positioning of the samples governed by cell type. B cells and PBMCs were separated from DCs and monocytes in PC2 (Figure 1B). Along PC1 pDC samples clustered together, followed by tDCs, cDC2, DC3 and then classical (mo1), intermediate (mo int) and non-classical monocytes (mo2). Timepoint after vaccination contributed little to the overall variance. Thus, circulating subsets of DCs and monocytes largely maintain their transcriptional identity after vaccination.

**Figure 1.**
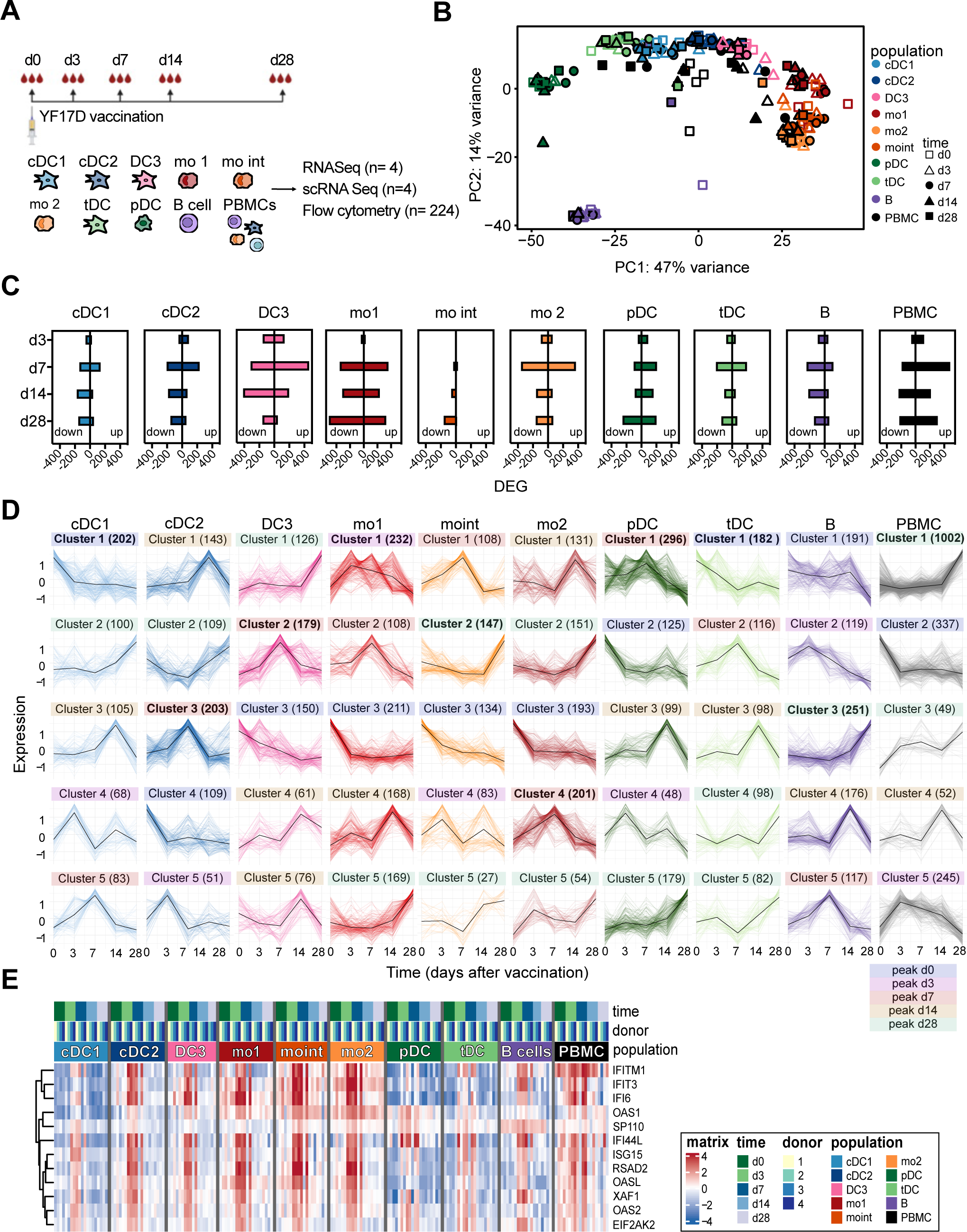
YF17D vaccination induces common and distinct time-dependent transcriptome changes in blood APC subpopulations. (A) Experimental design of the study. (B) PCA analysis of VST-transformed bulk RNA-seq data after pre-filtering. Sorted populations are indicated by colors and the symbols indicate the time after vaccination. (C) DESeq2 was used to find the significantly up- and downregulated genes (adjusted p < 0.05, Bonferroni corrected) of each population and each time point after YF17D vaccination compared to day 0. The number of up- and downregulated DEGs are shown and the colors indicate the populations. (D) Clustering of expressed genes (log counts per million ≥ 0.5) with significant changes in expression between time points (ANOVA, p <0.05) for each population over time. Colored by population. Number in brackets indicates genes in each cluster. Cluster colors indicate peak of gene expression within the cluster on day 0 (blue), day 3 (pink), day 7 (red), day 14 (orange) or day 28 (green). Clusters in bold have highest gene number compared to other clusters within the population. (E) Heatmap showing hierarchically clustered DEG upregulated on day 7 vs. day 0 common to all APC subsets (cDC1, cDC2, DC3, mo1, moint, mo2, tDC, pDC, B, PBMCs). Gene expression is indicated by the color (red: high expression, blue: low expression) for each donor and all DC and monocyte subpopulations, B cells and PBMCs. Timepoints, vaccinees and subpopulations indicated by colors above the heatmap.

Gene signatures were extracted that allow distinction of DC subpopulations after acute viral stimulation. Cell type defining genes preserved after vaccination included *CADM1*, *CLEC9A* and *XCR1* for cDC1; *CD1C*, *GRIP1* and *CD1E* for cDC2; *VSIG4*, *RNASE1* and *DTNA* for *DC3*; *PLEKHD1*, *SMIM5* and *VASH2* in pDCs; *AXL*, *SIGLEC6* and *ADAM33* in tDCs (supplemental Figure S1, Table S3). While some populations like cDC1, pDCs and tDCs showed clearly distinct gene expression profiles, cDC2, DC3 and monocytes gene signatures were overlapping, confirming the similarity of these populations seen in the PCA. DC3 characteristically expressed genes found in both monocytes (e.g. *F13A*) and in cDC2 (e.g. *CD1C*).

The greatest variance in gene expression was explained by cell type, but comparison of individual timepoints after vaccination to baseline within each population revealed time-dependent transcriptomic changes (Figure 1C, table S3). The highest average numbers of significantly up- and downregulated genes (differentially expressed genes, DEGs) were found on day 7 after vaccination in most cell types, indicating the peak of the systemic innate immune response. Both mo1 and DC3 showed a higher number of DEGs compared to the other populations, while mo int were the least responsive cell type. In mo1 and pDCs changes in transcriptome were still observed 28 days after vaccination, while gene expression in the other populations returned to baseline or showed only few DEGs at that time point.

By hierarchical clustering of genes showing significant changes in expression over time, we identified distinct temporal patterns (Figure 1D, Supplementary Table S4). Each cell population exhibited multiple clusters with different dynamics. In the majority of DC and monocyte subpopulations (cDC2, DC3, mo1, mo2, pDCs) the largest clusters (Figure 1D, indicated in bold) were those with a peak on day 7 and/or day 3 after vaccination confirming the results of the DEG analysis shown in Figure 1C. Notably, each cell population, except for PBMCs, had a cluster with peak gene expression on day 7 post-vaccination (Figure 1D, red). Overrepresentation analysis using MSigDB Hallmark and C2 canonical pathways showed significant enrichment for the Reactome pathway “Interferon gamma signaling” and “Interferon alpha response” or “Interferon alpha/beta signaling” in these predominant clusters across all populations (Supplementary Table S4).

In cDC1 and tDC, the largest clusters showed the highest level of gene expression before vaccination followed by a decrease on day 3 and fluctuation at later timepoints. In cDC1 cluster 1 genes from the Hallmark pathways “TNFα signaling via NF-κB” and “P53 pathway” were overrepresented. Conversely, tDCs cluster 1 showed no significant pathway enrichment but contained genes such as *RAD1*, *ZBTB40* involved in cell cycle checkpoints, and transcriptional regulation. In PBMCs, B cells, and moint the largest clusters emcompassed genes with increased expression on day 28 compared to the other time points: In B cells cluster 3 with this dynamic was enriched for genes from Reactome pathways “Cellular response to chemical stress”, “Fatty acid metabolism,” and KEGG pathway “Oxidative phosphorylation”. Within the PBMC cluster 1, we found genes such as *IFNLR1*, *GZMB* and *VEGFB* while in moint cluster 2 we found genes such as *HMGB1*. Temporal gene clusters with a similar upregulation at the later timepoints were also found in the other subsets most prominently in mo1 (cluster 5) and pDCs (cluster 5) consistent with the high number of DEGs observed between d28 and d0 in these subsets. In mo1 cluster 5 was enriched for ISGs, Hallmark “E2F targets”, “Adipogenesis” and “Fatty Acid Metabolism”. We also observed biphasic clusters with peaks on day 3 and day 14. These were enriched for Reactome “Interferon alpha/beta signaling” and “Interferon gamma signaling” in mo2 and Hallmark “Fatty acid metabolism” and “Adipogenesis” in mo1. Thus, common and diverse dynamics of gene expression were observed in the different populations highlighting the complex and coordinated early immune response to YF17D vaccination.

Comparing the DEGs of day 7 vs baseline between the different APC subsets we identified a group of commonly upregulated genes (Figure 1E). This shared response gene signature consisted of 12 interferon stimulated genes (ISGs): *IFITM1*, *IFIT3*, *IFI6*, *OASL*, *SP110* (also known as *IFI41* or *IFI75*), *IFI44L*, *ISG15*, *RSAD2*, *OASL*, *XAF*, *OAS2*, *EIF2AK2*. Depending on the individual vaccinee these genes were upregulated on day 3 and/or on day 7 and returned to baseline on day 14 after vaccination reflecting a regulated temporary ISG response. Thus, distinct populations of APCs showed common and distinct dynamics of gene expression and shared a core ISG response peaking on day 3 or 7 after yellow fever vaccination.

Gene set enrichment analysis (GSEA) comparing each time point with the baseline before vaccination in all APC populations using all expressed genes confirmed the common enrichment of ISGs on days 3 and 7 after vaccination (Figure 2A). Reactome “Antigen processing and presentation” was enriched in all DC subsets, in mo int and mo2 at all time points with peak on day 7 (Figure 2A and supplementary Figure 2B). Gene sets involved in translational pathways and oxidative phosphorylation were underrepresented in mo1 and pDCs on days 3, 7, and 14 and in cDC1 on day 3, but were enriched in cDC2, DC3 and tDCs at all timepoints and cDC1 at later timepoints indicating differences between cell types in the metabolic response and impact of the vaccination on ribosome biogenesis and translation (Figure 2A).

**Figure 2.**
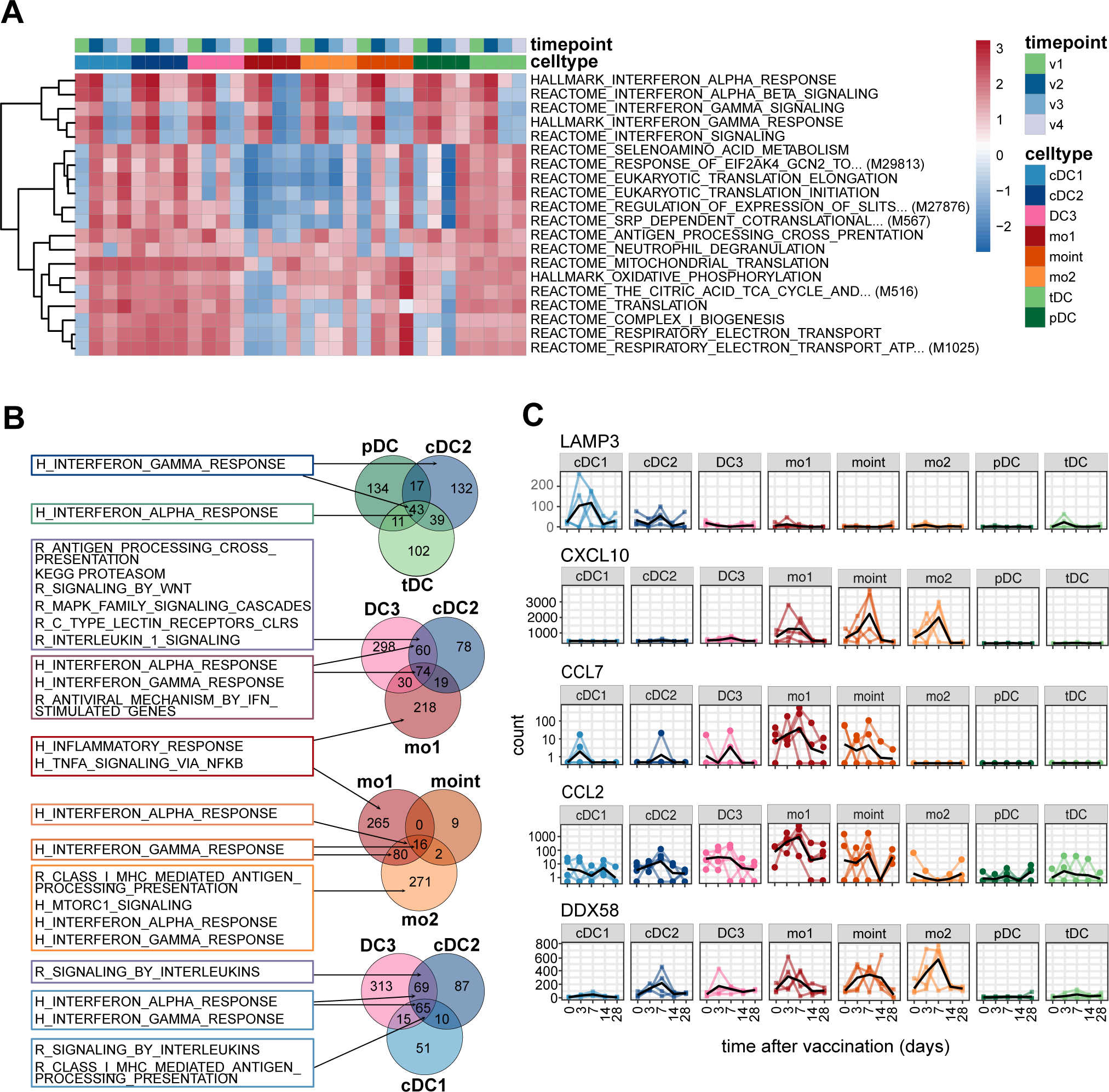
Transcriptome response of circulating APC subsets to YF17D is dominated by ISGs. (A) Heatmap of unscaled normalized enrichment scores (NES) for selected pathways from HALLMARK and REACTOME across four time point comparisons against baseline. Displayed pathways include only those with NES greater than 2 in at least one comparison. Gene set enrichment analysis (GSEA) was applied using the t-statistic of each comparison. Upper margin colors indicate populations and timepoints. The color scale bar represents NES values. (B) Venn diagrams showing the number of overlapping and distinct DEGs upregulated on d7 vs. d0 (padj. < 0.05) in the indicated set of populations. Hallmark (H) and Reactome (R) pathways identified by overrepresentation analysis are shown for DEGs unique to one population or overlapping between populations for each Venn diagram. (C) Normalized counts in different populations of the indicated genes over time after vaccination. Values for individual donors are connected by colored lines. Black lines indicate the mean.

We next compared DEGs (d7 vs d0) between related APC populations. tDCs shared more response genes with cDC2 than with pDCs. Hallmark gene set “Interferon gamma response” was overrepresented uniquely in cDC2 when compared to pDCs and tDCs. DC3, mo1, and cDC2 had many overlapping peak response genes with overrepresentation of ISGs as expected. Response genes shared between DC3 and cDC2 were enriched for “antigen processing and cross-presentation”, “proteasome”, “signaling by Wnt”, “MAPK family signaling cascades”, “C-type lectin like receptors and “Interleukin 1 family signaling” while response genes unique for mo1 were enriched in Hallmark “inflammatory reponse” and “TNF-α signaling via NFκB”. This unique response in mo1 was also observed in comparison with mo int and mo2. Unique among the monocyte subsets mo2 response genes showed overrepresentation of “Class I MHC mediated antigen processing and presentation”. cDC1, cDC2, and DC3 shared many peak response genes. Besides ISG these were enriched for Reactome gene sets “Class I MHC mediated antigen processing and presentation” and “signaling by interleukins” highlighting their common antigen presentation function.

Further analysis of cell-type specific response genes revealed for example upregulation in cDC1 of lysosomal associated membrane protein 3 (*LAMP3*) implicated in DC maturation and MHC class II presentation (de Saint-Vis et al., 1998) and transient upregulation of chemokine expression, e.g. *CXCL10*, *CCL2* and *CCL7* particularly in monocytes (Figure 2C). *DDX58*, encoding viral RNA sensor RIG-I was also significantly upregulated in monocytes, and less so in cDC2 and DC3 suggesting that these cell types become more responsive to RIG-I ligands after vaccination (Figure 2C).

Thus, besides a common ISG-dominated response to YF17D we detected cell type specific changes in gene expression suggesting distinct functional contributions of different DC and monocyte subpopulations.

### Single cell RNA Seq reveals activated cell clusters expressing ISGs and inflammatory genes on day 3 and 7 after vaccination

The expression of response genes in defined cell populations may be heterogeneous with several differentiation and activation states coexisting within the same subpopulation. To discover distinct activation states within APC subpopulations in an unbiased manner, we performed single-cell RNA sequencing (scRNA-seq) of the DC and monocyte fractions sorted from PBMCs of 4 donors (2 males and 2 females) before and at day 3 and 7 after vaccination. Sorted DCs and monocytes of each donor were combined at a ratio of 4:1 to enrich for rare DC subsets. After removing doublets and cells with high mitochondrial RNA (>7.5%) or low total RNA (<200 genes) we analyzed ∼35,000 cells. Individual donors where integrated using harmony normalization. Unbiased clustering revealed 14 clusters with clusters 3, 4, 6, 7 found mostly on days 3 and 7 after vaccination (Figure 3A, Supplemental Figure S3B). These clusters showed high expression of ISGs exemplified by *IFI44L* and *ISG15* indicating an activated antiviral cell state (Figure 3B, 3C).

**Figure 3.**
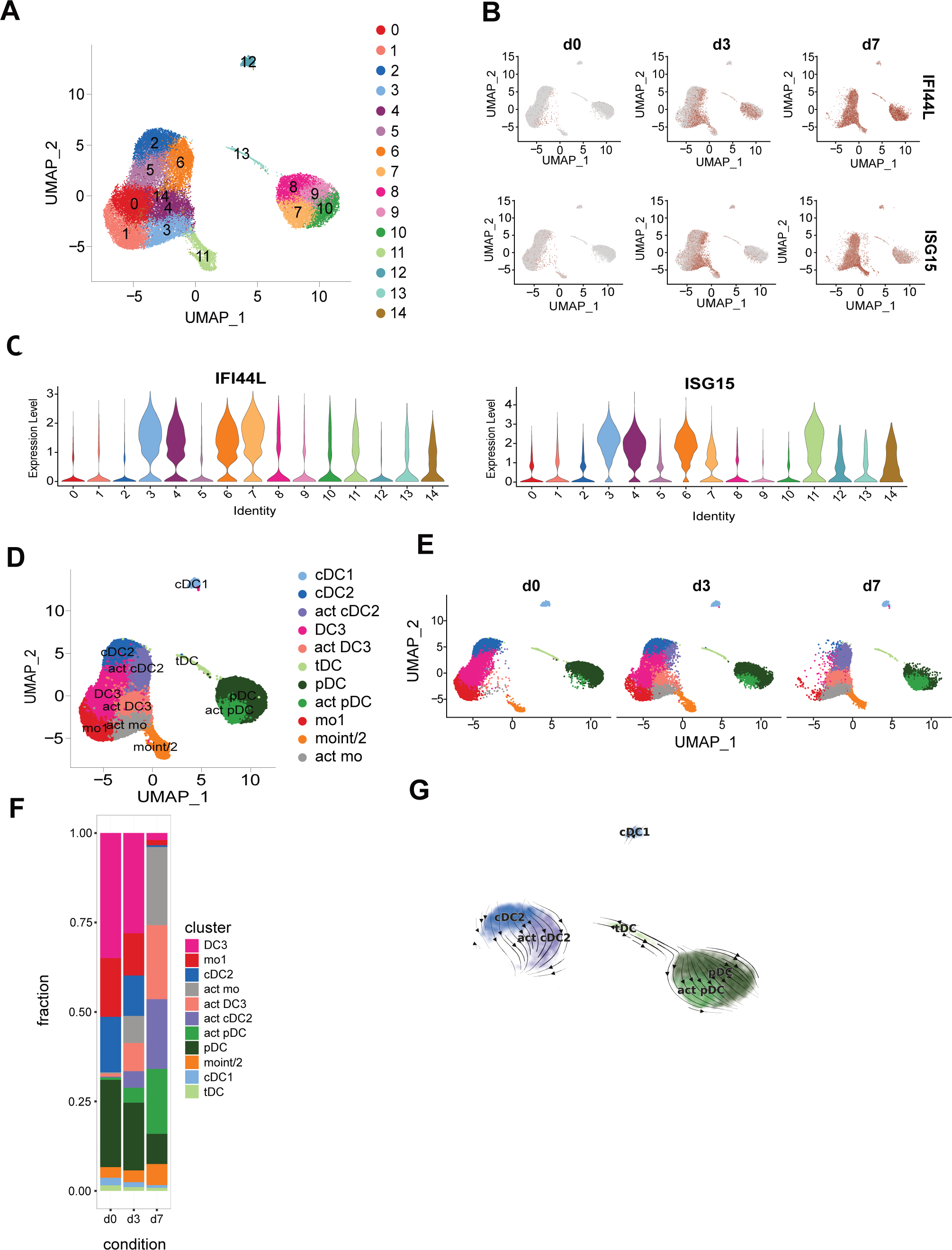
scRNA-seq indicates collective acquisition of an activated cell state in blood APCs early after vaccination. (A) UMAP visualization of all scRNA-seq data of sorted HLA DR^+^ Lin^-^ (CD3, CD19, CD20, CD56) living cells generated using the 10x Genomics workflow after cleanup and harmony normalization. Louvain clusters based on gene expression are shown in different colors and are numbered. (B) UMAP visualization from A showing expression of IFI44L and ISG15 as color overlay. Red = high expression, grey = low expression. (C) Violin plots showing expression of key functional markers ISG15 and IFI44L in the individual clusters. (D) UMAP visualization with annotated cell clusters shown as colored overlay (act: activated, indicated by high ISG expression). (E) UMAP visualization of scRNA-seq data from different time points (before (d0) and at d3 and d7 after vaccination) with the annotated cell clusters indicated by colors as in D. (F) Stacked bars show the frequency of the individual populations annotated before (d0) and at d3 and d7 after vaccination. (G) The RNA velocity vector field derived from scvelo analysis of cells identified as cDCs, pDCs and tDCs (all donors and timepoints combined) was projected onto the UMAP with annotated cell clusters indicated by colors.

The individual clusters were annotated manually using known marker genes for DC and monocyte subpopulations derived from publications (Villani, Satija et al. 2017, Dutertre, Becht et al. 2019) and our own bulk RNA gene signatures (Supplementary Figure S3A and S3D). After merging highly similar clusters, cDC1, cDC2, DC3, mo1, mo int/mo2, tDC and pDC populations as well as activated cDC2, activated DC3, activated pDC and activated monocytes were identified (Figure 3D). The percentages of cells found in the activated DC and monocyte clusters greatly increased on day 3 and 7. This ISG-driven response was most prominent in cDC2, DC3 and mo1 where on day 7 the activated clusters almost entirely replaced the clusters found on day 0 (Figure 3E and 3F). In 3 donors a partial shift towards more activated cells was already seen on day 3 after vaccination (Supplementary Figure 3C). These results demonstrate that most cells responded to the vaccination in a concerted fashion especially cDCs and monocytes with full activation reached by day 7. *IFN-Typ I/II* transcripts were not detected above threshold in this data set at any of the time points investigated (Figure 4C) similar to the bulk RNA-seq results (supplementary Figure 2). IFN-α2, IFN-β, IFN-γ, IFN-λ1 and -λ2 were measured in plasma samples from day 0, 3, 7, 14 and 28 in a subgroup of 22 vaccinees using a bead-based multiplex assay, but could not be detected in any of the samples (unpublished data).

**Figure 4.**
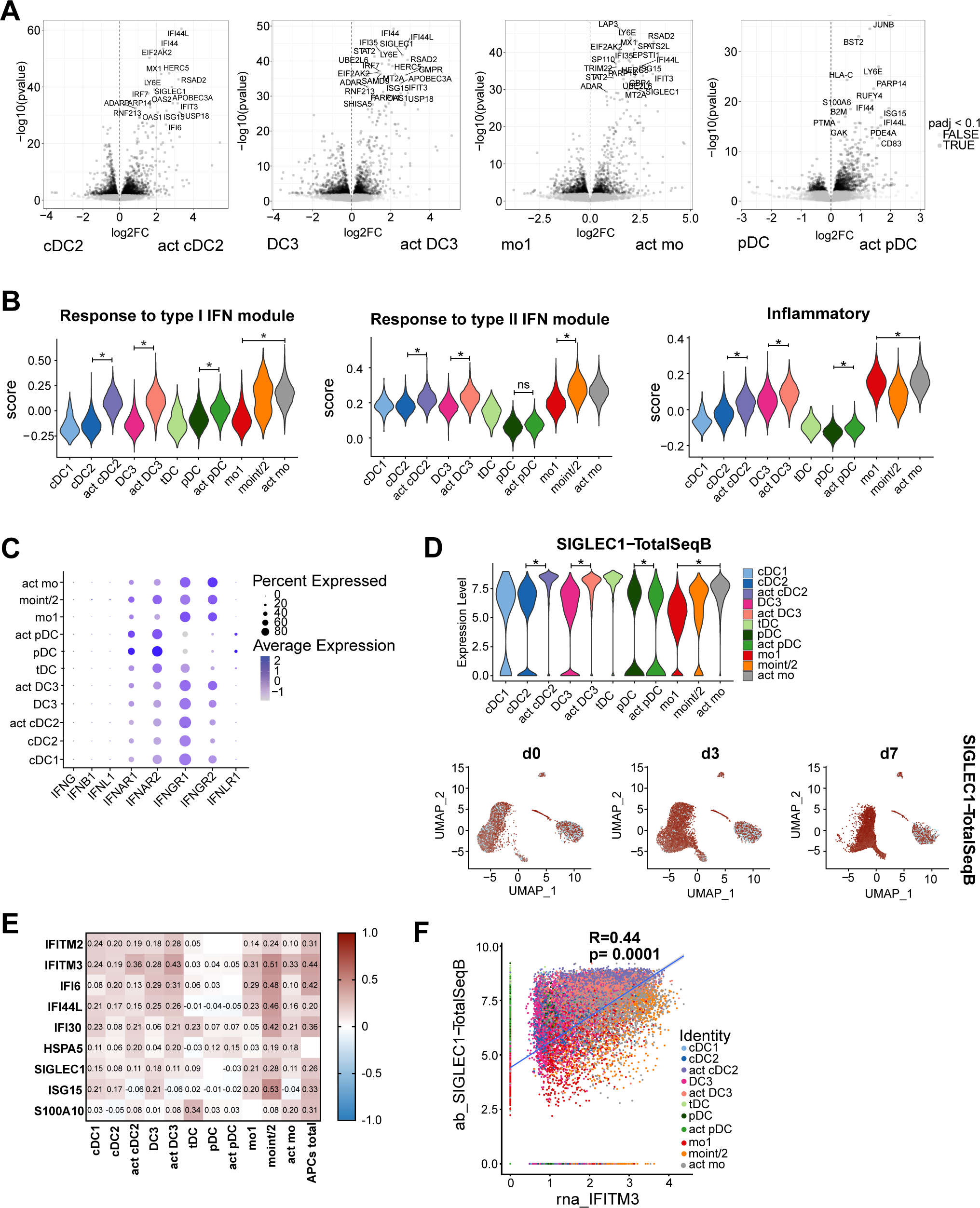
SIGLEC1 expression on the cell surface marks the IFN-induced activation state in cDC2, DC3 and monocytes. (A) Volcano plots showing comparison of activated and non-activated clusters of cDC2, DC3, mo1 and pDCs with annotation of the most significantly up-and downregulated (by p-value, log2 fold-change > 0.3) genes. (B) Violin plots showing the indicated gene expression scores based on average counts of the module genes in each cell cluster. Significant difference of scores between activated and non-activated cell clusters are indicated by asterisks (Wilcoxon-test using the mean module scores of the individual donors, n=4). (C) Dot plot showing the expression of all detected IFN and IFN receptor genes in the cell clusters. Color intensities indicate the average gene expression values and diameters indicate the percentages of cells expressing the genes within the cluster. (D) Violin plot showing surface expression of SIGLEC1 as measured using TotalSeqB antibodies. UMAP showing surface expression of SIGLEC1 detected by TotalSeqB antibody as color overlay projected onto the separate UMAP before and on day 3 and 7 after vaccination. Red: high expression; grey: low expression. (E) Pearson correlation analysis of surface SIGLEC1 expression. Correlation coefficients (R values) are shown in the heatmap for selected genes highly correlating with SIGLEC1 gene expression in each population. (F) Scatter plot of IFITM3 RNA expression and SIGLEC1 expression of all cells with linear regression line. Colors indicate belonging of cells to a specific cell cluster. Pearson correlation coefficient R and p-value are indicated in the graph.

Since tDCs were previously shown to differentiate into cells with a cDC2 phenotype in vitro (See, Dutertre et al. 2017) and in mice in vivo (Rodrigues et al., 2023; Sulczewski et al., 2023) we inferred trajectories from RNA velocity analysis of the combined data of cDC and pDC subsets from all time points using the scVelo method. RNA velocity vectors were bidirectional pointing from tDCs towards cDC2 as well as pDCs (Figure 3G). These results suggest that human tDCs are heterogenous and contain cells with divergent trajectories towards pDCs and cDC2.

For cDC1 and tDCs where a separate activation cluster was not detected, we found significant upregulation of multiple ISGs on day 3 and 7 after vaccination compared to baseline confirming the bulk RNA-seq results (Supplemental Figure 4). Pairwise comparison between the activated and nonactivated APC clusters confirmed upregulation of ISGs such as *IFI44L*, *EIF2AK2* and *ISG15* in activated clusters of cDC2, DC3, monocytes and pDCs (Figure 4A). Relative expression scores derived from the type I IFN induced gene module (Edahiro et al., 2023) were significantly higher in the activated clusters (Figure 4B and Table S6). The type II IFN induced gene module scores were also significantly increased in the activated compared to the non-activated cDC2 and DC3 clusters and in the mo int/mo2 cluster compared to mo1 (Figure 4B). This was not the case for the activated pDC cluster coinciding with a lower mRNA expression level of IFN-γ receptors *IFNGR1* and *IFNGR2* in pDCs compared to cDCs and monocytes (Figure 4C). Relative expression scores of the hallmark inflammatory gene set also tended to be increased in the activated compared to the non-activated cDC2 and DC3 clusters and were constitutively high in monocytes and low in pDCs and tDCs (Figure 4B). For cDC1 and tDCs where a separate activation cluster was not detected, we found significant upregulation of multiple ISGs on days 3 and 7 after vaccination compared to baseline confirming the bulk RNA-seq results (Supplemental Figure 4).

Expression of SIGLEC1 encoding for the CD169 surface molecule, which was one of the genes highly upregulated in the activated cDC2, DC3 and mo1 clusters (Figure 4A), was quantified on the protein level in the same experiment using TotalSeq B antibody labeling and sequencing. It was highly expressed in tDCs and in the activated cDC2, DC3 and mo1 clusters (Figure 4D). Surface expression of SIGLEC1 protein correlated positively with mRNA expression of different ISGs such as *IFITM2*, *IFITM3*, *IFI6*, *IFI30* and *ISG15* within all cells analyzed together and in each individual cluster except for tDCs and pDCs (Figure 4E, F). Therefore, surface expression of SIGLEC1 is a sensitive marker for the early response to IFNs and marks the temporary activated state of circulating cDCs and monocytes after YF17D vaccination.

### Upregulation of SIGLEC1 coincides with upregulation of costimulatory molecules and activation markers on the surface of cDCs and monocytes

To further characterize the functional state of circulating DC and monocyte subsets before and after YF17D vaccination and investigate interindividual variation of this response, we analyzed the phenotype of DC and monocyte subsets in cryopreserved PBMC samples by multi-dimensional spectral flow cytometry in a larger number of vaccinees. To link phenotype and transcriptome results, SIGLEC1/CD169 was included as a marker in addition to costimulatory molecules and chemokine receptors. First, PBMC from all 4 timepoints were analyzed in a subgroup of 10 patients. We detected temporary upregulation of mean fluorescence intensities (MFI) for CD86, PD-L1 and SIGLEC1 on day 7 after vaccination in all populations except for tDCs and pDCs consistent with the transcriptomic data. SIGLEC1 was already upregulated on day 3 and further increased on day 7 (Figure 5A and 5C). Expression of AXL, another IFN induced surface molecule, was induced on day 7 in all DC subsets and in mo1 (Figure 5A). CD83 was upregulated in cDC2, DC3 and monocytes on day 7. CD40 was also upregulated in many subsets on day 7 except in cDC1 and DC3. HLA-DR was upregulated in monocytes but not in DCs. CXCR3 expression was induced whereas CCR2 expression was reduced on day 7 in mo1 and in pDCs, whereas CCR2 expression was induced in mo2 indicating differential regulation of chemokine receptor expression between cell types after the vaccination. Population frequencies were also significantly altered after vaccination, with a shift from mo1 to moint and mo2 on day 7 and a slight reduction of cDC1 and cDC2 subsets (significant for cDC1) on day 7 after vaccination (Supplementary Figure S5A) confirming previous results obtained from freshly isolated PBMC (Winheim et al., 2023).

**Figure 5.**
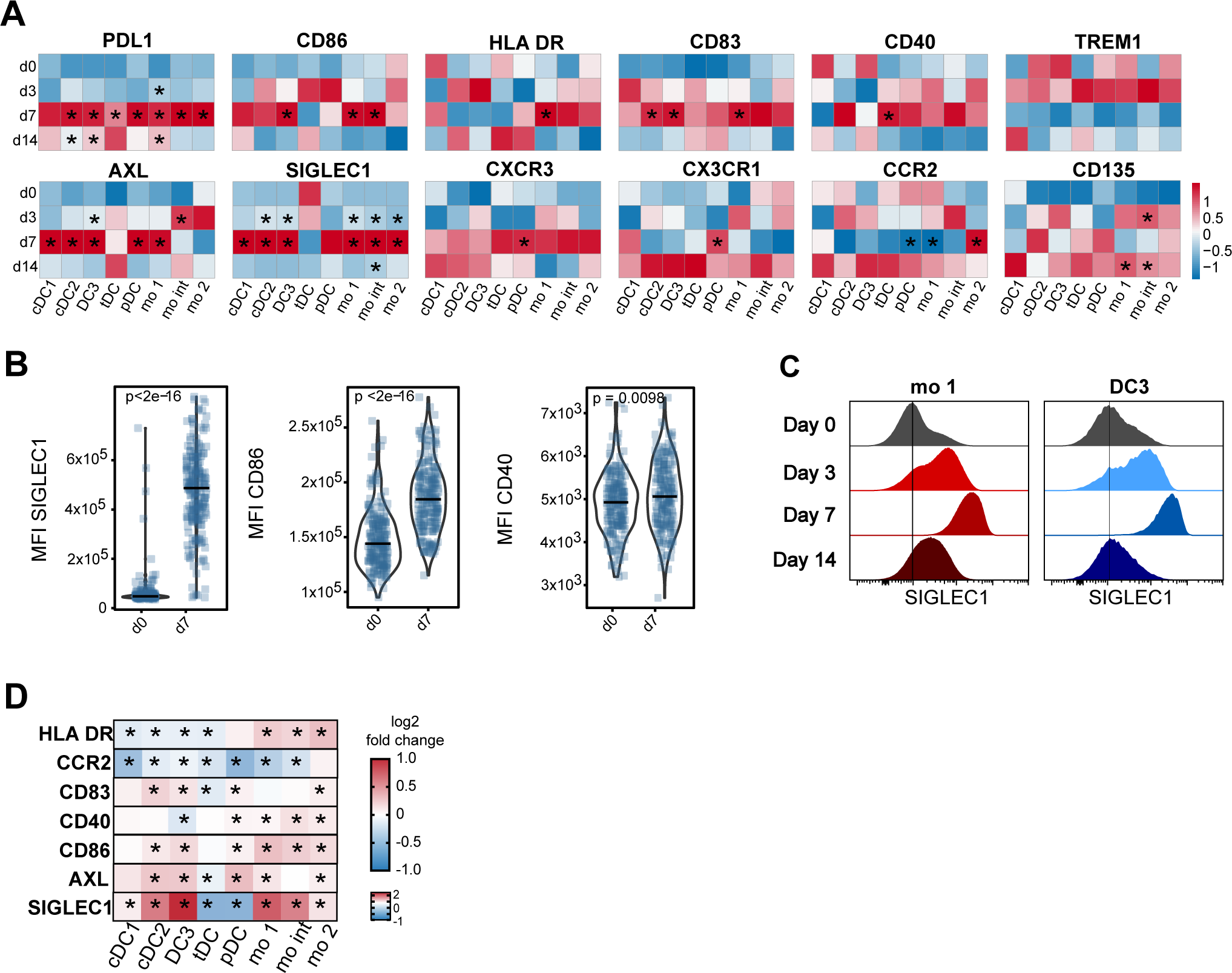
Temporary upregulation of SIGLEC1 and costimulatory molecules characterizes the response of circulating APCs to YF17D in a large cohort of vaccinees. Phenotype of blood DC and monocytes subpopulations analysed by flow cytometry. (A) Heatmap of the mean MFI values of indicated cell surface markers before vaccination (d0) and on days 3, 7, and 14 after vaccination (n=10). The data is scaled for each column and significant differences to baseline are indicated by asterisks (* p < 0.05). Red: high expression, blue: low expression. (B) Violin plots showing the MFI of SIGLEC1, CD86 and CD40 on mo1 on days 0 and 7 after vaccination (n=214). Measured values of individual study participants are indicated by blue-colored dots and p values shown in the graphs. (C) Histograms showing the SIGLEC1 expression level in mo1 and DC3 before vaccination (d0) and on days 3, 7, and 14 after vaccination (concatenated data of 10 donors). (D) Heatmap of the mean log2 fold change of the MFIs (d7 vs. d0) of the indicated surface markers (n=214). Red: positive log2fc; blue: negative log2fc. Significant differences to baseline are indicated by asterisks (* p < 0.05, Wilcoxon test, Bonferroni corrected).

We selected the timepoints day 0 and day 7 to perform the same analysis for a cohort of additional 214 vaccinees. Significant increases in the MFIs of CD86, CD83 and AXL were detected on day 7 in all DC and monocytes subsets except for tDCs (Figure 5B and 5D). Upregulation of SIGLEC1 was detected in all subsets, except tDCs and pDCs. CD40 was significantly upregulated in pDCs and monocytes, but not in cDCs. Downregulation of CCR2 was observed in all populations, except for mo2. HLA-DR was significantly upregulated in monocyte subsets but rather downregulated in cDC subsets (Figure 5D).

To identify novel activation clusters that might be missed by manual gating analysis, unbiased FlowSOM clustering was performed on the HLA DR^+^ Lin^–^ viable cells containing the DC and monocyte fractions. The clusters were visualized in a UMAP and assigned to monocytes or DCs according to presence or absence of CD88/CD89 expression (Supplementary Figure 5B, C, D). The DC and monocyte fractions were then each clustered separately. Clusters were annotated and similar clusters were fused (Supplementary Figure S5E). Within the DC compartment cDC1, cDC2, CD163^–^ DC3, CD163^+^ CD14^+^ DC3, SIGLEC1^+^ DC3, tDC, pDC were identified (Figure 6A and 6B). Monocyte clusters were annotated as mo1 (CD163^–^ SIGLEC1^–^), SIGLEC1^+^ mo1, CD163^+^ SIGLEC1^–^ mo1, CD163^+^ SIGLEC1^+^ mo1, mo int, mo2 (Figure 6C and 6D). Within the DC compartment cDC1 and cDC2 were reduced and pDCs slightly increased. In the monocytes the frequency of mo int was significantly increased as seen previously in the exploration cohort (Figure 6E). The frequency of cells in the SIGLEC1^+^ DC3 and monocyte clusters increased dramatically on day 7 and only few vaccinees had SIGLEC1^+^ cells at baseline with frequencies below 25% (Figure 6E). Thus, the upregulation of SIGLEC1 surface expression on day 7 after YF17D vaccination and the appearance of distinct SIGLEC1^+^ DC and monocyte clusters reflecting an IFN-induced activation state could be validated in a large cohort. Although an increase in SIGLEC1 expression was seen in the great majority of vaccinees on day 7 after vaccination, this response was variable between individuals warranting further investigation as a predictor of vaccination outcome.

**Figure 6.**
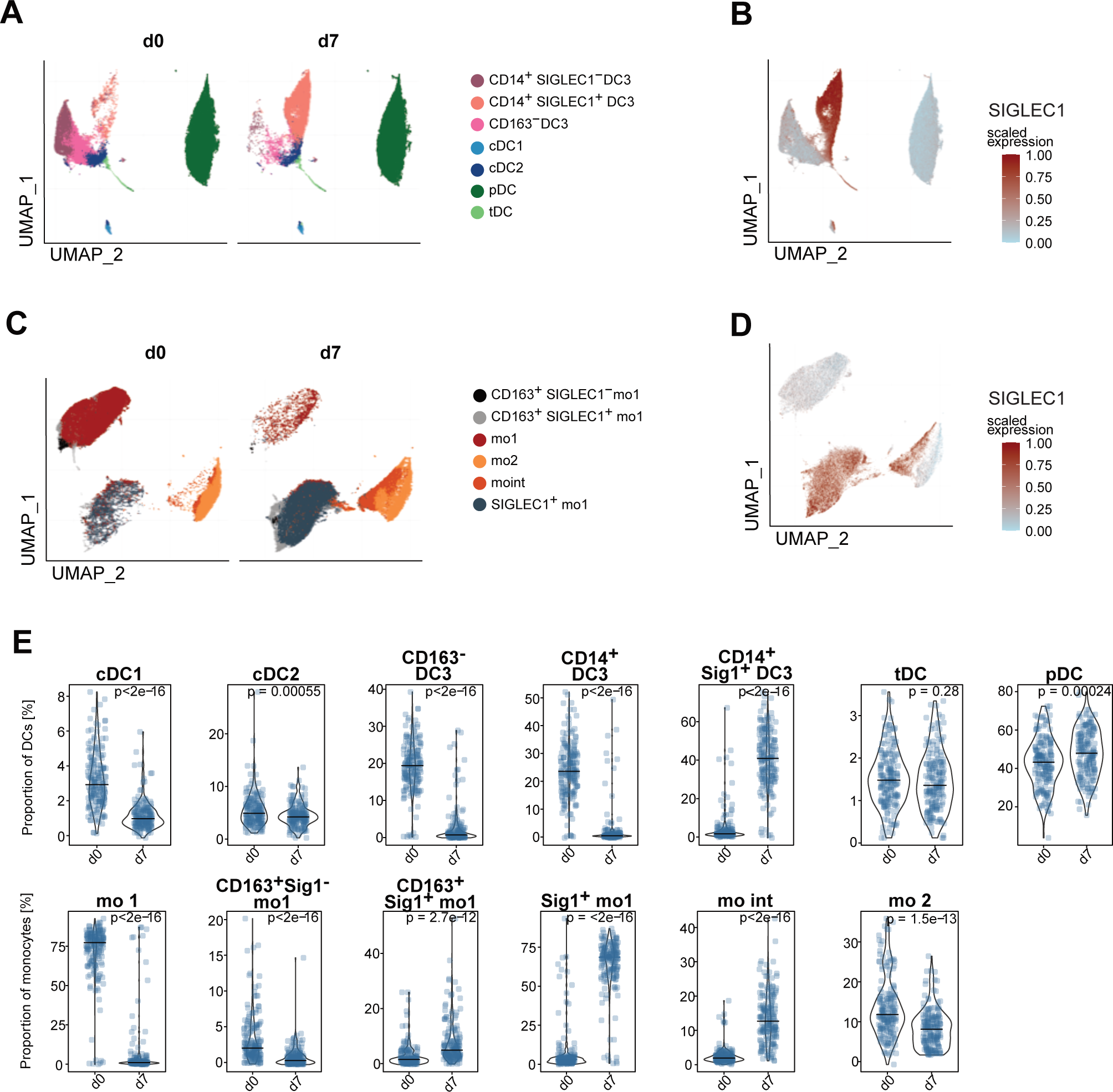
Unbiased clustering of flow cytometric data confirms generation of SIGLEC1-positive DC3 and monocyte clusters with interindividual variability. FlowSOM clustering was performed on the HLA-DR^+^ Lin^–^ living cells containing both the DC and monocyte fractions. The clusters were visualized in a UMAP and assigned to monocytes or DCs according to presence or absence of CD88/CD89 expression. Reclustering of DC and monocyte fractions was performed using FlowSOM. After annotating and fusing similar clusters the data was depicted in a UMAP with annotated clusters shown as colored overlay. (A and C) Annotated DC clusters (A) and monocytes clusters (C) on d0 and d7 after vaccination. (B and D) SIGLEC1 scaled expression indicated by red-brown color overlayed on the UMAP embeddings of DCs (B) and monocytes (D). (E) Violin plots show DC and monocyte subpopulation frequencies identified using FlowSOM clustering on d0 and 7 after vaccination as proportions of total DCs or total monocytes (n=214). Horizontal lines indicate the median. Measured values of individual study participants are indicated by blue-colored dots and p values shown in the graphs (Wilcoxon test, Bonferroni corrected).

### High SIGLEC1 upregulation on cDCs and monocytes is associated with high protective antibody titers early after YF17D vaccination

Vaccination with YF17D induced high titers of neutralizing antibodies (dominated by IgM isotype) on day 14 and 28 after vaccination indicating protection against infection was reached in all vaccinees of our cohort with highly variable titers (Santos-Peral et al., 2024). We hypothesized that the IFN-induced activation of blood cDCs and monocytes indicated by upregulation of SIGLEC1 surface expression relates to the rapid generation of protective antibody and T cell responses characteristic for YF vaccination. To explore this, we categorized individuals based on high and low SIGLEC1 fold-change (d7 vs d0) in cDC1, cDC2, DC3, mo1, and mo2 through unsupervised clustering (Figure 7A). pDCs and tDCs were excluded from the analysis due to the absence of SIGLEC1 upregulation post vaccination (Fig 4 and Fig 5). Interestingly, the extent of SIGLEC1 upregulation was consistent across the different APC populations suggesting a concerted action and association with vaccine responsiveness. Stratification of vaccinees into high and low SIGLEC1 upregulation, with the exclusion of intermediate levels, allows a precise examination of this signature’s impact on subsequent vaccine responses, which exhibit high variability across individuals.

**Figure 7.**
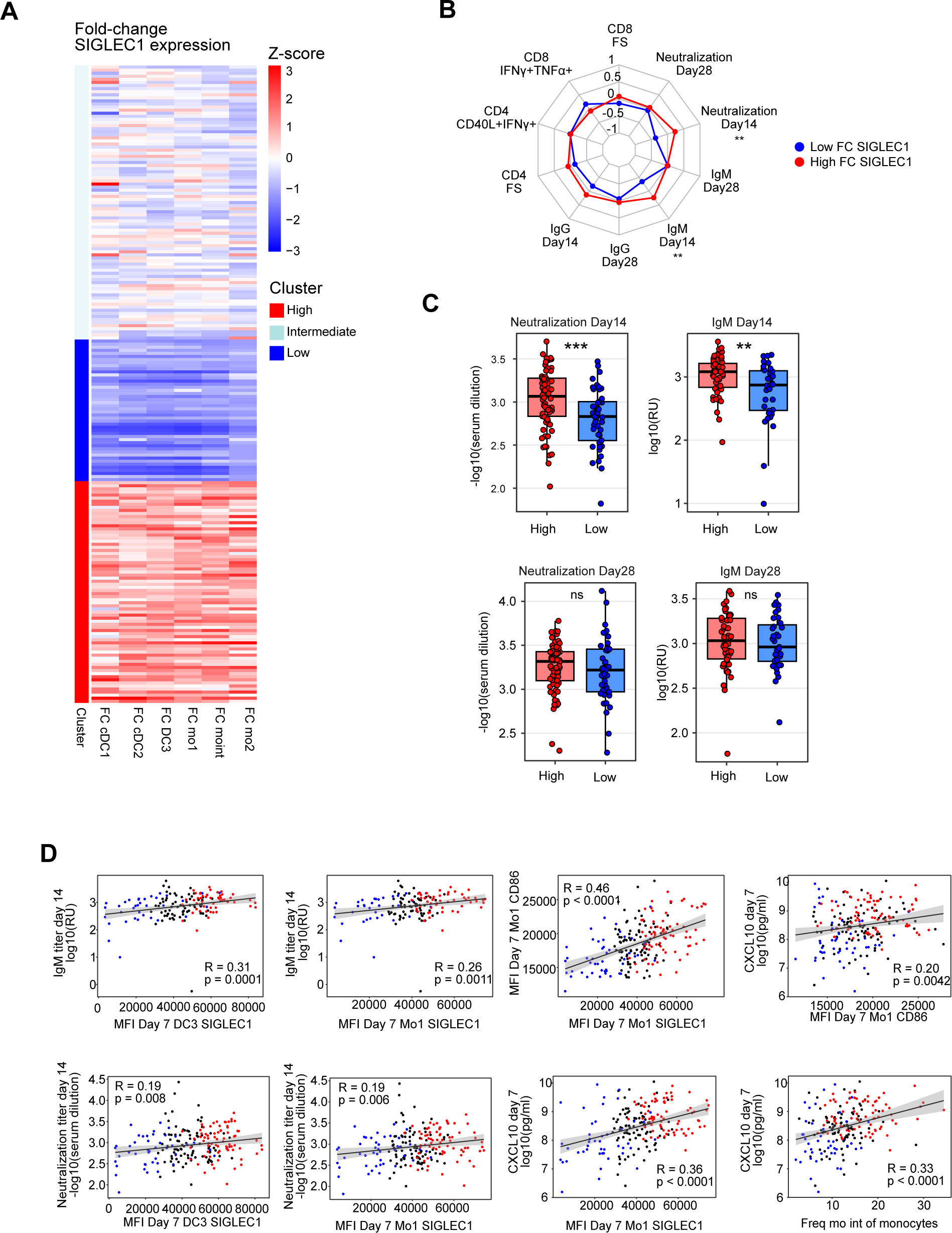
SIGLEC1 upregulation is associated with the early protective antibody response to YF17D. A) Unsupervised hybrid hierarchical k-means clustering performed on the scaled fold-changes between day 7 and baseline in the expression of SIGLEC1 for DC and monocyte populations. (B) A radar plot illustrating the association of vaccination endpoints for the two clusters identified in A, denoting high SIGLEC1 upregulation in red and low in blue. Vaccination endpoints include antigen-specific CD4 and CD8 functional scores (FS), the frequency of antigen-specific CD40L^+^ IFN-γ^+^ CD4^+^ T cells, the frequency of antigen-specific IFN-γ^+^ TNF-α^+^ CD8^+^ T cells and the yellow fever-specific IgG, IgM, and neutralizing antibody titers on day 14 and day 28 post-vaccination. The data was scaled, and the radar plot depicts the means, with significance estimated using a Student’s t-test. (C) Boxplots comparing the neutralization and IgM antibody titers on day 14 and 28 for individuals with high or low SIGLEC1 upregulation identified in A. Significance was calculated using a Wilcoxon rank-sum test. (D) Spearman correlation analysis was conducted between the indicated parameters. Correlation coefficients, p-values, and linear regression lines are shown for all individuals and for those with high (red) and low (blue) SIGLEC1 upregulation identified in A. Statistical significance is depicted as follows: *pC≤C0.05, **pC≤C0.01, ***pC≤C0.001 for B and C.

Vaccinees with high SIGLEC1 upregulation on day 7 showed significantly higher neutralizing antibody titers and virus-specific IgM titers early after vaccination (day 14) than vaccinees with low SIGLEC1 upregulation (Figure 7B-C). Antigen-specific T cell responses and antibody responses on day 28 were not significantly altered between the subgroups (Figure 7B-C). Spearman rank correlation analyses confirmed the correlation of SIGLEC1 MFI on day 7 and fold change (d7 vs d0) in DC3 and mo1 with early antibody titers. SIGLEC1 and CD86 expression correlated with each other and with day 7 plasma CXCL10 concentrations representing the systemic IFN-induced cytokine response. The frequency of mo int on day 7 also positively correlated with plasma CXCL10 levels at the same time point. In sum, the temporary IFN-induced activation of circulating APCs detected by SIGLEC1 upregulation on day 7 after YF17D vaccination is associated with the early protective antibody response.

## Discussion

Our study provides a comprehensive analysis of the temporal dynamics of gene expression in individual blood DC and monocytes subpopulations following YF17D vaccination in humans. We observed a transient ISG response within the first week after vaccination as the predominant feature in all subsets and defined a common set of significantly upregulated ISGs playing a crucial role in the antiviral immune responses. Specifically, *IFI44L*, *OAS2*, *OAS3*, *IFIT2*, *IFIT3*, and *RSAD2* have been shown to impede viruses by disrupting viral replication and translation (Busse et al., 2020; Kimura et al., 2013; Schoggins et al., 2011; Teng et al., 2012; Terenzi et al., 2006). Additionally, members of the IFN-inducible transmembrane family, such as *IFITM1* and *IFITM3*, obstruct viral entry and hinder early stages of the life cycle of various viruses (Wrensch et al., 2015). Consequently, this robust ISG signature reflects a systemic antiviral state within the circulating DC and monocyte compartment after YF17D vaccination which contrasts with the downregulation of ISGs that is seen in severe Dengue and Zika virus infections (Ghita et al., 2023; Sun et al., 2017). In the context of YF17D vaccination, several studies have identified the importance of a functional type I IFN response to control replication and pathogenicity of the vaccine virus. Patients with IFNAR1 and IFNAR2 deficiency or with autoantibodies against multiple type I interferons can present with life-threatening complications after vaccination (Bastard et al., 2021; Hernandez et al., 2019). Therefore, the antiviral state caused by upregulation of these ISGs seems to be highly relevant for controlling YF17D and ensuring the safety of this live vaccine.

Direct contact between the virus and APCs could induce the ISG response observed here as the virus is present in the blood with peak viremia between days 3 to 9 after vaccination (Akondy et al., 2015). DCs can be infected and respond to direct contact with the YF17D virus through the activation of diverse pattern-recognition receptors, triggering cellular activation and cytokine secretion, including IFN-α (Barba-Spaeth et al., 2005; Bruni et al., 2015; Cong et al., 2016; Douam et al., 2017; Querec et al., 2006). However, expression of IFN transcripts in the various APC subsets in the peripheral blood was minimal or undetectable (in the case of IFN-α) by bulk and scRNA-seq, in the individuals and time points used for this study. In line with our findings, Hou et al. failed to detect IFN-α/-β/-λ mRNA expression in PBMC at earlier time points (4 and 24 hrs) in a cohort of 21 vaccinees (Hou et al., 2017). Additionally, our scRNA-seq data revealed a synchronized shift of the majority of cells within DC and monocyte subpopulations toward ISG-expressing clusters after vaccination consistent with a response to IFNs produced at the vaccination site or other tissues containing the vaccine virus and released into the circulation. However, neither IFN-α/-β nor IFN-γ or IFN-λ could be detected in plasma samples of our cohort using a sensitive bead-based multiplex assay. But even in acute flares of systemic lupus erythematosus with high ISG expression in PBMCs IFNs are only detectable in the plasma using ultrasensitive methods (Mathian et al., 2019) suggesting that IFNs could be present in the blood of our vaccinees at very low concentrations. We observed a higher expression of type I IFN module genes compared to type II IFN module genes in blood APCs suggesting that type I IFNs are the dominant cytokines inducing the ISG response, but IFN-γ produced early after YF17D vaccination by NK cells (Neves et al., 2009; Neves et al., 2013) may also contribute. It was described for SARS-CoV-2 mRNA vaccination for example that the frequency of myeloid cells with increased ISG expression emerging shortly after booster vaccination correlated with plasma IFN-γ levels at this timepoint (Arunachalam et al., 2021).

SIGLEC1/CD169 expression is induced by IFN-α, IFN-β, and IFN-ω, but not IFN-γ (Bourgoin et al., 2020) highlighting its role in antiviral defense (Herzog et al., 2022) and as a marker of acute viral infection (Bedin et al., 2021; Sakumura et al., 2023; Winheim et al., 2021). We identified SIGLEC1 upregulation as a sensitive indicator of the early systemic response to IFNs and the transient activation state elicited in blood cDCs and monocytes by YF17D vaccination, which correlated with plasma CXCL10 levels and early antibody responses. A transient induction of SIGLEC1/CD169 expression on monocytes and CD1c^+^ DCs associated with the IFN response signature was also reported after booster vaccination with a VSV-vectored HIV-1 vaccine (Hao et al., 2021). In our study simultaneous upregulation of SIGLEC1 and costimulatory molecules indicated a temporary IFN-mediated functional maturation of cDCs as a characteristic feature of YF17D vaccination that may promote induction of adaptive immunity. We found an association of high SIGLEC1 upregulation on day 7 with high neutralizing IgM antibody titers on day 14 after vaccination providing a link between the systemic IFN response and rapid production of high titers of protective antibodies. Our findings resonate with an Ebola vaccine study that demonstrated upregulation of ISGs by the rVSVΔG-ZEBOV-GP vaccine and correlation with subsequent protective antibody titers (Vianello et al., 2022). The type I IFN response may be directly linked to rapid antibody induction via activity of type I IFN on B cells promoting differentiation into antibody secreting cells (Jego et al., 2003; Swanson et al., 2010). Alternatively, IFN-induced activation of APCs could support B cell differentiation in germinal centers and antibody response via induction of T follicular helper cells (Dahlgren et al., 2022), which we found to be rapidly activated within the first week after YF17D vaccination correlating with neutralizing antibody titers (Huber et al., 2020). Gressier et al showed that conditioning of murine DCs with type I IFNs increased responsiveness to CD40 engagement by CD40-L expressed on T helper cells leading to optimal priming of CD8^+^ T cells (Gressier et al., 2023). However, we did not find a positive correlation of SIGLEC1 expression in DCs or monocytes with the frequency of YF17D-specific CD8^+^ T cells on day 28 after YF17D vaccination suggesting that the positive effect of APC activation may be counteracted by other mechanisms. For example, IFN-mediated reduction of viral replication may limit the availability of viral antigen for CD8^+^ T cell induction, which was shown to be determined by the YF17D viral load (Akondy et al., 2015).

While all blood APC populations exhibited upregulation of a common set of ISGs and activation markers on day 7 post-vaccination, distinct subset-specific responses to YF17D vaccination were evident underscoring the diverse and specialized functions of APC subpopulations in response to vaccination. pDCs for example did not significantly upregulate SIGLEC1 but other ISGs suggesting a differential response to IFNs and acquisition of a different activation state than cDCs and monocytes. The specific upregulation of LAMP3 in cDC1 may indicate preferential conversion of this cell type to a mature migratory DC type which has been described in tumors and inflamed tissues (Nakamizo et al., 2021; Zhang et al., 2019). Monocytes and DC3 showed heightened expression of various chemokines, including CXCL10, CCL7 and CCL2, aligning with their pro-inflammatory potential (Dutertre et al., 2019). The similar transcriptomic responses of cDC2, DC3 and monocytes after vaccination with YF17D are consistent with the reported convergence of transcriptomic changes in DC3, monocytes and cDC2 following type I IFN stimulation demonstrating their close relationship and distinction from other APC subtypes (Girard, Law et al. 2020).

In conclusion, our study presents a comprehensive analysis of the transcriptomic responses of blood APC subpopulations to YF17D vaccination, highlighting a global ISG response within and across diverse subsets as well as cell-type specific responses. Notably, we found that upregulation of SIGLEC1 expression which represents IFN-induced activation in blood cDCs and monocytes is associated with early protective antibody titers, unveiling a pivotal link between IFN-mediated APC activation and humoral immunity. Thus, our study provides novel insights and resources to inform development of vaccines to rapidly induce protective immunity in naïve individuals.

## Material and Methods

### Sample Collection and study design

The yellow fever vaccination study was conducted by the Department of Clinical Pharmacology of the University Hospital, LMU Munich, Germany. The study protocol was approved by the Institutional Review Board of the Medical Faculty of LMU Munich (IRB #86-16) and adhered to the most recent version of the declaration of Helsinki. The study cohort was registered in the ISRCTN registry (ISRCTN17974967). All participants were healthy (aged 19 to 44 years, Table S1) and had not been previously exposed to wild-type YFV, and were not previously immunized against YF. After giving informed consent, the patients received a single subcutaneous injection of the YF17D vaccine (Stamaril, Sanofi Pasteur, Lyon, France) at the Division of Infectious Diseases and Tropical Medicine at LMU Munich. Blood was drawn directly before vaccination and on days 3, 7, 14, and 28 after vaccination using the S-Monovette Sodium-Heparin (Sarstedt, Nürnbrecht, Germany). Peripheral blood mononuclear cells (PBMCs) were isolated by Ficoll density gradient centrifugation and frozen in 90% heat-inactivated FCS/10% DMSO (v/v) in liquid nitrogen. Plasma samples were collected in Sodium EDTA monovettes (Sarstedt, Germany) centrifuged and frozen in liquid nitrogen.

### Flow cytometric analysis of DC and monocyte subpopulations

Cryopreserved PBMC samples of vaccinees containing 1.5-3 x 10^6^ cells were thawed, processed, stained, and analyzed by flow cytometry in four batches. The exploratory batch of 10 donors included timepoints day 0, 3, 7 and 14. The follow-up cohort of 214 patients included only day 0 and day 7 after vaccination and was separated into 3 batches measured within one week of 60, 82, and 78 vaccinees each.

PBMC were stained in 50μl of PBS, 2mM EDTA, 10% FCS (v/v) containing FcR blocking reagent (Miltenyi Biotec) with fluorescently labeled antibodies as indicated in Table S3 and incubated for 30 min at 4°C and washed in PBS, 2mM EDTA, 10% FCS (v/v) three times. Fixable viability dyes were used according to the manufacturers’ protocol. Cells were fixed with BD Cytofix (Cat. # 554655), washed three times and resuspended in PBS, 2mM EDTA, 10% FCS (v/v). Samples were measured using the Cytek Aurora (Cytec Biosciences) with the recommended Cytek assay settings, where gains are automatically adjusted after each daily QC based on laser and detector performance to an optimal value, ensuring comparability between measurements. Unmixing was performed using single stained cells or beads.

Analysis of flow cytometric data was performed in Flow Jo, version 10.8.1. Samples were unmixed at the Cytek Aurora and compensation adjusted in Flow Jo. Viable single cells in the HLA DR^+^ Lin^–^ gate were exported from Flow Jo and used for further analysis in R with flowWorkspace::open_flowjo_xml (flowWorkspace version 4.2.0). Cells with negative FI and failing upper boundary filtering were removed. Data was converted to a SingleCellExperiment using CATALYST::prepData (version 1.14.0) with parameters FACS = T and cofactor = 150 for arcsine transformation.

FlowSOM clustering was performed separately on three batches that were stained and measured on individual days within the same week. Samples were clustered and separated into monocytes and DCs based on expression of CD88/CD89 Rphenograph (version 0.99.1). Then DCs and monocytes were reclustered separately and annotated. Data was visualized using CATALYST functions. Cluster frequencies and MFIs were exported and batch correction was performed using first BoxCox transformation with fpp (v 0.5) and combat batch correction from sva (v 3.18).

### Cell sorting

For sorting APC populations for bulk RNA sequencing analysis, T cells were separated via CD3 magnetic bead isolation, and the CD3 negative cell fraction was stained with antibody master mix, washed and subsequently used for cell sorting. Cells were sorted using an 85 µm nozzle at a BD FACSAria Fusion (BD Biosciences either directly into RLT lysis buffer for B cells or into RPMI 1640 (Biochrom, 10% FCS, 100 U/ml penicillin, 100 μg/ml streptomycin, 1% non-essential amino acids, 1 mM sodium pyruvate, 2 mM GlutaMAX, 0.05 mM β-mercaptoethanol, 2 mM EDTA) and then resorted directly into RLT lysis buffer. Cells were then vortexed, centrifuged (450 g, 4°C, 5 min), and then immediately stored at -80°C until further processing. For sorting HLA-DR^+^ Lin-cells for scRNA sequencing analysis, PBMCs were directly stained with antibody master mix, washed and used for cell sorting. Cells were sorted using a 100µm nozzle and sorted into PBS, 2mM EDTA, spinned down immediately and resuspended in PBS, counted using a Neubauer chamber and used for 10x 3′ single-cell capture (Chromium Next GEM Single Cell 3’ Kit v3.1).

### Bulk sequencing using SmartSeqv2

RNA was isolated from sorted cells using the Qiagen RNeasy Plus Microkit (Qiagen, Hilden, Germany) according to the manufacturer’s instructions. The RNA was eluted in 10 µl DEPC water and the RNA quality was determined using the Bioanalyzer RNA 6000 pico assay (Agilent Technologies, Santa Clara, California, USA) according to the manufacturer’s instructions. The Smart-Seq2 protocol was performed on 3.7 µl of isolated RNA with 0.2 µl of 1:125,000 diluted ERCC spike-ins as described previously (Picelli, Faridani et al. 2014) with minor modifications. Briefly, PCR preamplification products were purified using Ampure XP beads in a 0.6:1 ratio, tagmentation was done using ILMN Tag DNA Enzyme & Buffer Small Kit (Illumina # 20034197) decreasing tagmentation reaction volume (to 2.5μl) (Beltran, Gerdes et al. 2019). PCR purification was carried out using Ampure XP beads in a 0.75:1 ratio. Samples were then sequenced with NextSeq1000 using 100 bp paired-end sequencing by the LAFUGA Genzentrum RNA sequencing data was demultiplexed from FASTq files using the Illumina demultiplex tool of the LAFUGA Gene Center (Munich), that was custom-written by Alexander Graf and allows one mismatch in the i5/i7 sequence. The adapter sequences were removed, and the reads were filtered for gene length. The quality of the reads was controlled with FastQC (v0.11.7), and reads were aligned to the human genome (UCSC hg38) using the RNA STAR seq mapper (v2.7.8a). Four samples were excluded due to insufficient read quality (donor 1: B cells d14, DC3 d7 and d2; donor 3: mo int d28). The number of reads was counted using HTSeq-count (0.6.1p1) and differential gene expression analysis was performed using DESeq2 (v1.34.0) on all cell populations to first identify unique population identifying genes that are not affected by vaccine response (Supplementary Figure S1). DESeq2 was also used on all cell populations together to identify common gene expression changes driven by timepoint after vaccination, as well as on each cell population seperately to identify population specific effects driven by timepoint after vaccination. The data was transformed with variance stabilizing transformation (VST) for PCA visualization and filtered for genes < 50 counts. For clustering of genes with similar time course in each cell population (Figure 1D) we first excluded genes with zero counts in all samples and then genes with log-counts per million < 0.5. Genes with significant expression changes over time (ANOVA, p-value < 0.05) were retained for clustering. The pairwise Euclidian distances between genes were calculated using the dist() function of R package stats (v 4.3.0). Genes were hierarchically clustered based on the distance matrix to group genes with similar expression patterns over time. The cutree function was used and the optimal number of clusters was determined using the elbow method. GSEA analysis (Figure 2A) was performed using fgsea and ranked gene lists. Venn diagramms were plotted using VennDiagramm (v 1.7.3). The enricher function from the clusterProfiler package was used for over-representation analysis (ORA) against the MSigDB Hallmark (H) and C2 canonical pathways. Multiple testing corrections were applied using the Benjamini-Hochberg method to control the false discovery rate (FDR). Only pathways with adjusted p-values (FDR < 0.05) were considered to be significantly overrepresented. Annotation of the Venn diagrams was restricted to overrepresented pathways with count ≥ 10 and gene ratio ≥ 0.15.

### Single cell RNA sequencing using 10x Chromium

Gene expression libraries were prepared according to the manufacturer’s protocol (10x Genomics). All libraries were sequenced using a NovaSeq 6000 (Illumina) to achieve 25,000 reads/cell for gene expression and 5,000 reads/cell for protein expression. Droplet libraries were processed using Cell Ranger v4.0. Reads were aligned to the GRCh38. Data was analyzed using Seurat. Raw data was filtered to remove cells with fewer than 200 genes and with high mitochondrial RNA > 7.5%. Each cell was given a doublet score using DoubletFinder and doublets were removed by excluding cells with multiple or no hashtags. Donor bias was removed using Harmony normalization. Data were log-normalized and highly variable genes were identified using the Seurat vst algorithm. Louvain clustering was performed on the first 20 PCs and initial resolution of 1.1. Proliferation score was calculated but not affecting clustering and therefore not regressed. Clusters were annotated using published gene signature sets (Villani, Satija et al. 2017, Leylek, Alcantara-Hernandez et al. 2019) as well as gene signature sets derived from our bulk RNA sequencing data and then merged. DEGs were determined using DESeq2 after aggregating reads per cell type per donor (n=4). Donor ID was included as random factor in the statistical model. Type I and type II IFN response module scores were calculated as described (Edahiro et al., 2023) using the gene ontology gene sets Response to Type I IFN (<u>GO:0034340</u>) and Response to Interferon Gamma (<u>GO:0034341</u>), respectively.

For RNA velocity analysis in scVelo data was exported from Seurat and, using loompy, combined with loom files obtained by velocyto processing of cellranger alignment files. Data was filtered for tDC, cDC2, activated cDC2, cDC1, pDC, and activated pDC. We calculated moments with ‘scv.pp.moments’ on 30 principal components. Splicing kinetics were obtained with ‘scv.tl.recover_dynamics’, followed by calculating velocities (‘scv.tl.velocity’, mode=’dynamical’) and constructing a velocity graph (‘scv.tl.velocity_graph’). Finally, we visualized the results using ‘scv.pl.velocity_embedding_stream’, using UMAP embedding, colored by cell type.

### Cytokine detection in plasma samples

Cytokine concentrations were measured in plasma samples using three panels (pro Human Cytokine, pro Human Chemokine, pro Human Inflammation) of the Bio-Plex Multiplex Assay (Bio-Rad Laboratories, USA) following the manufacturer’s instructions. The data were then analyzed using the Bio-Plex Manager Software and the concentrations of cytokines were log-transformed. The full panels were measured using plasma samples from 22 vaccinees from all timepoints. 18 selected analytes including CXCL10 were measured in the whole cohort before and on day 3 and 7 after vaccination.

### Quantification of anti-YF17D antibody titers in serum of YF17D vaccinees

The neutralizing antibody titers in serum samples were quantified by a Fluorescence Reduction Neutralization Test (FluoRNT) as previously described (Scheck et al., 2022). Briefly, the frequency of YF17D-Venus virus-infected Vero cells in the absence of vaccinee serum was set as 100% and the percentage of reduction was calculated for serial serum dilution steps. Neutralization curves were fitted by 4-parameter logistic regression using Prism 8 (GraphPad, La Jolla, CA, USA). 50 and 80 % FluoRNT values were interpolated from the curves. YF17D-specific antibodies in human serum samples were quantified with an in-house ELISA assay using recombinantly produced soluble E protein or YF17D-virion as antigens as described (Santos-Peral et al., 2023). These results have been published in a separate manuscript (Santos-Peral et al., 2023).

### T cell restimulation with YF17D virus and intracellular cytokine staining

To measure antigen-specific T cell responses, cryopreserved PBMC were thawed and rested overnight at high density (5.10^6^ million cells per mL) in R10 medium at 37°C in a 5% CO2 humidified atmosphere. PBMC were stimulated with live YF17D virus (1.5 10^7^ PFU/mL) or with the equivalent volume of purified supernatant of uninfected cells (unstimulated control) for 20 hours. Brefeldin A (BioLegend) and anti-CD107a (clone H4A3, BD Biosciences) were added for the last 4 hours of stimulation. Staining for intracellular cytokines and CD40L and flow cytometric analysis was performed as described (Santos-Peral et al., submitted manuscript). The data was analyzed in R with COMPASSimple package with a minimum of 100.000 iterations and 8 repetitions.

### Statistical analysis

Statistical analysis for scRNA seq data was performed as described above. Statistical analysis of flow cytometry data was performed either GraphPad Prism (v 9.1.0) or in R using ggpubr (v 0.5.0) kruskal wallis test, or wilcoxon test with Bonferroni correction as indicated in the individual figure legends.

### Clustering of vaccinees and correlation analysis

Vaccinees were clustered according to the fold-change in SIGLEC-1 expression in DCs and monocytes using hybrid hierarchical k-means clustering. Data were scaled and clustered using the hkmeans function from the factoextra R package using default settings. The optimal number of clusters was determined by the NbClust function (index = all). Hierarchical clustering was performed using Ward’s minimum variance method, applying the Euclidean distance measure to normalized data, along with the Hartigan-Wong algorithm for K-means. Spearman rank correlation analysis was performed in R.

## Acknowledgements

The authors thank Yvonne Schäfer for technical assistance.

The authors also thank Arne Kroidl, Günter Fröschl and Kristina Huber for serving as clinical study investigators. We acknowledge the Core Facility Flow Cytometry of the Biomedical Center, LMU Munich, and thank Lisa Richter and Pardis Khosravani. We acknowledge Eduardo Beltran for providing valuable advice. The authors also thank all the cohort participants who voluntarily participated in the study and donated samples. Parts of this work have been performed for the doctoral theses of EW, ASP, MZ at the LMU Munich.

## Author contributions

EW designed and performed experiments, analyzed and interpreted data, and wrote the manuscript; ASP, LR and MZ performed experiments, analyzed and interpreted the data; TE and SR analyzed and interpreted data and wrote the manuscript; MP helped to initiate the cohort, performed vaccinations, served as clinical study investigator, acquired funding, and supervised a part of the project. KE contributed to experimental design; TS contributed to experimental design, analyzed and interpreted data; GBS contributed unique reagents and acquired funding; SRo set up the cohort, contributed to experimental design, interpreted data, acquired funding and supervised; ABK conceived the study, designed experiments, interpreted data, acquired funding, supervised and wrote the manuscript. All authors critically reviewed the manuscript and approved the final version.

## Data availability

RNA-sequencing data is available in a public data repository. DOI accession number All other data is available upon request

## Funding

This work was funded by the Deutsche Forschungsgemeinschaft (DFG) project no. 210592381 SFB1054-TPA06 to A.B.K., project no. 369799452 TRR237-TPB14 to A.B.K. and S.R. The project received additional support from FlavImmunity a combined grant of the DFG project no. 391217598 to SR and ABK and the French National Research Agency (ANR) project no. ANR-17-CE15-0031-01 to G.B.S. The project received further funding by the European Union (YELLOW4FLAVI) project no. 101137459 to S.R., A.B.K. and G.B.S., and by grants of the iMed consortium of the German Helmholtz Societies and the Einheit für Klinische Pharmakologie (EKLIP), Helmholtz Zentrum München, Neuherberg, Germany to S.R. E.W. received funding from the Friedrich-Baur-Stiftung and a scholarship from the Villigst Foundation.

## Notes

### Competing Interest Statement

The authors have declared no competing interest.

