## Supplementary figures and images for "Interferon-induced activation state of circulating dendritic cells and monocytes triggered by yellow fever vaccination correlates with early protective antibody responses"

### Supplementary Figures

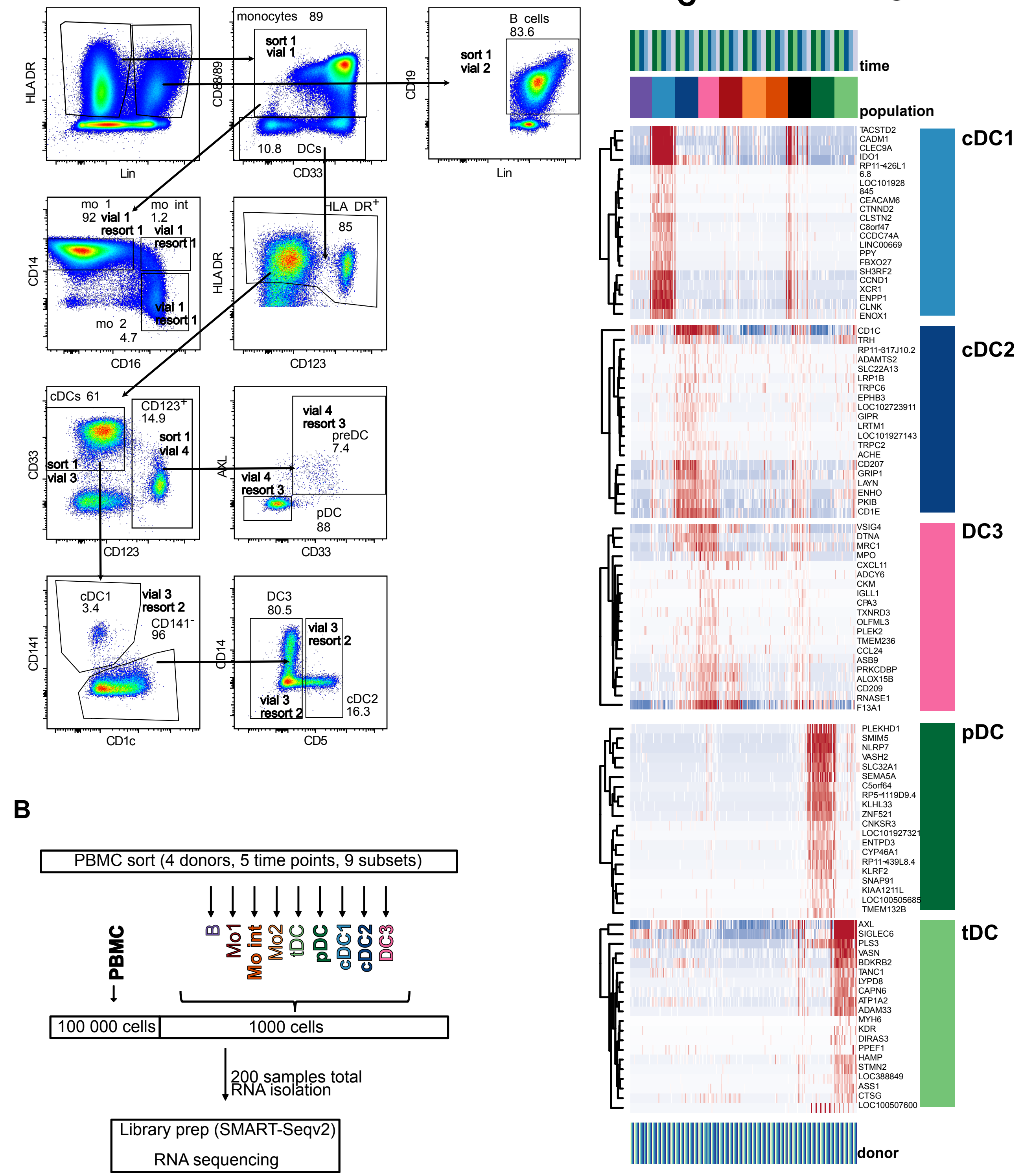

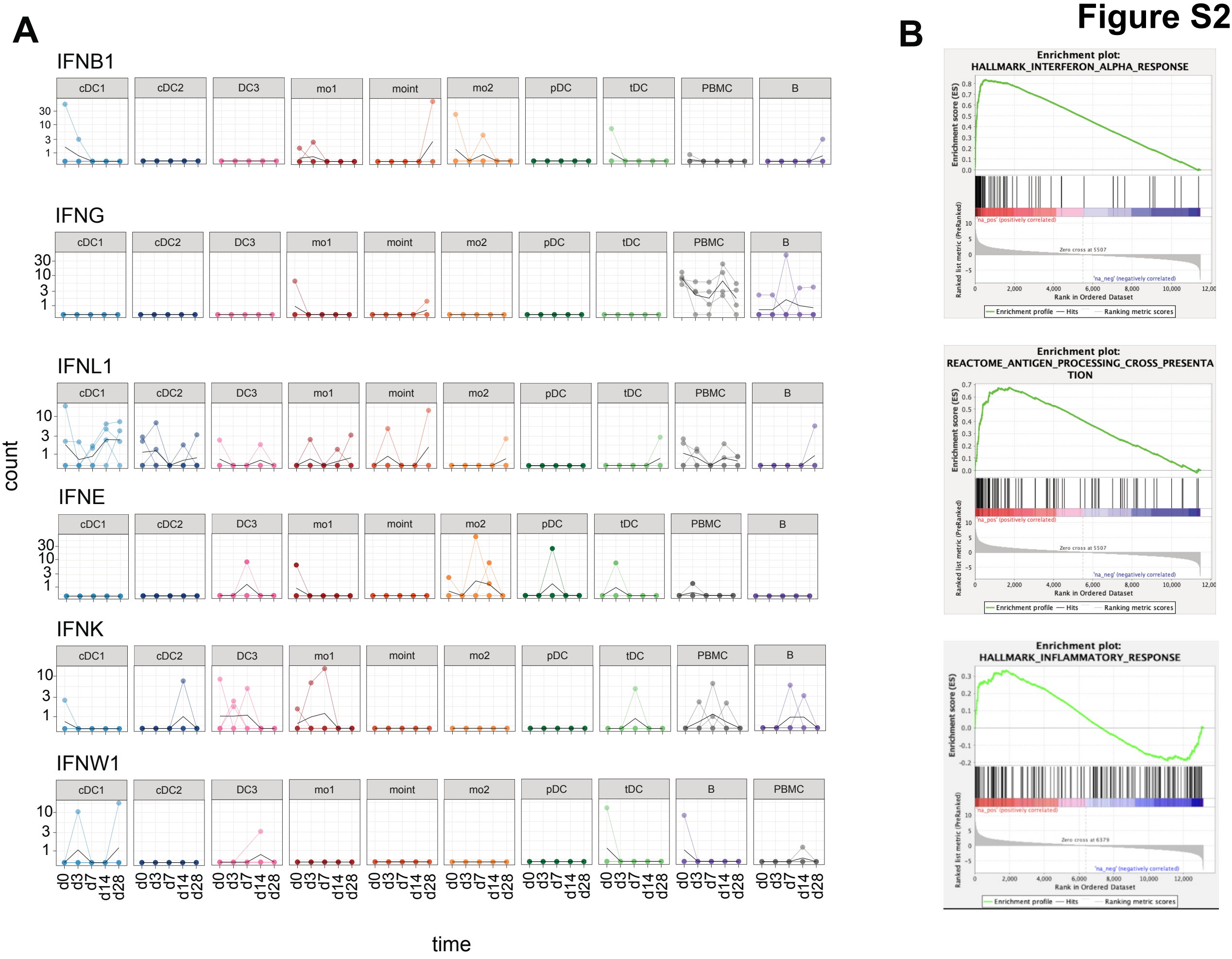

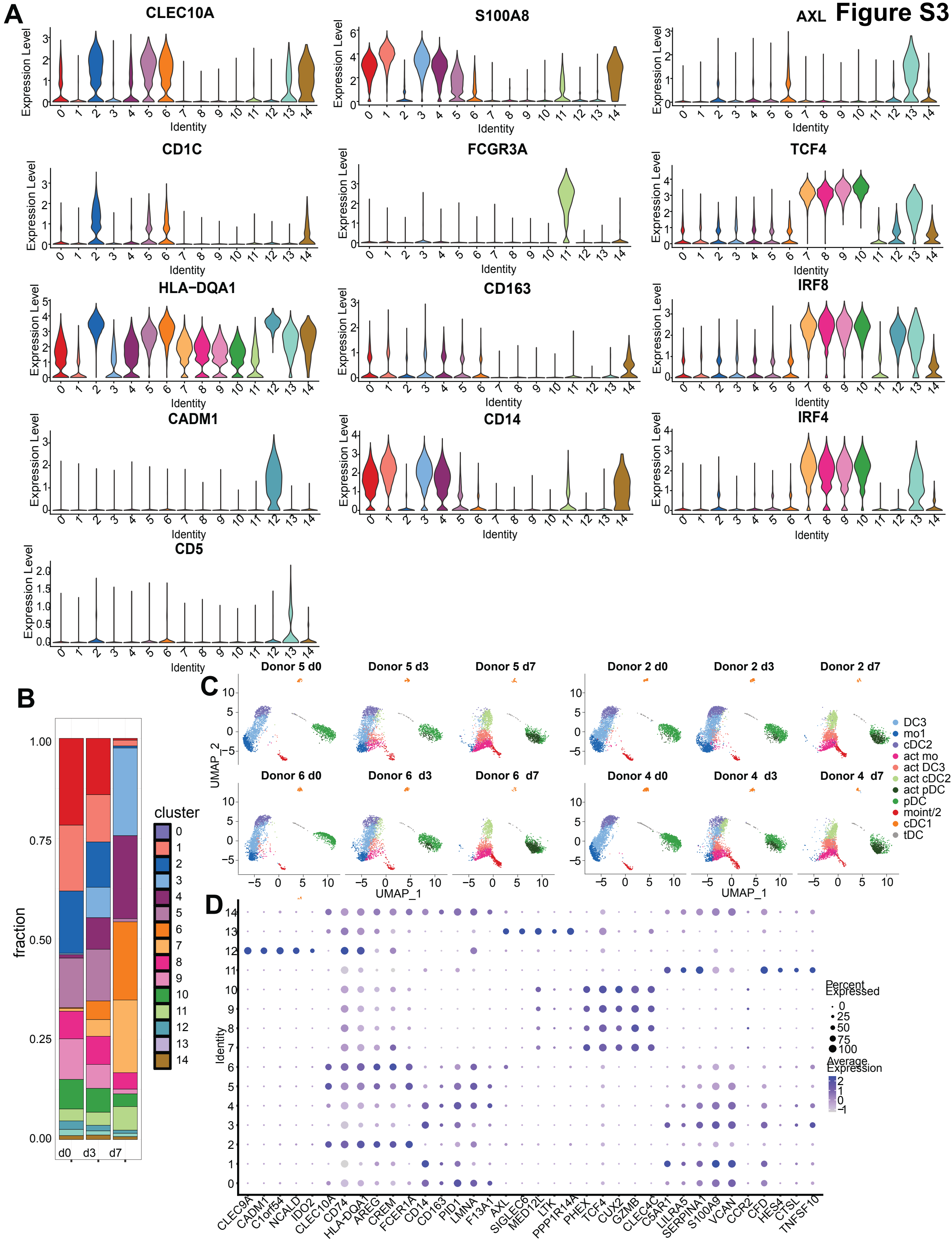

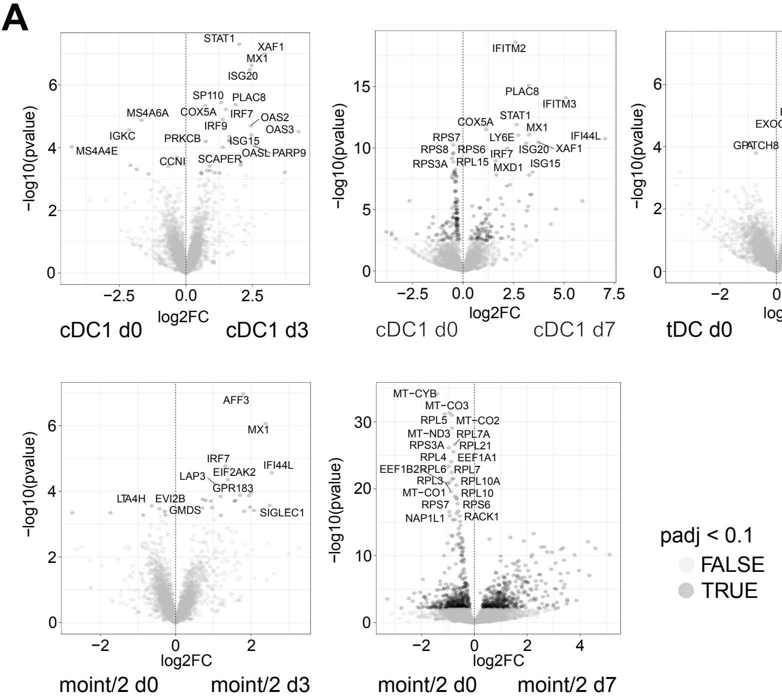

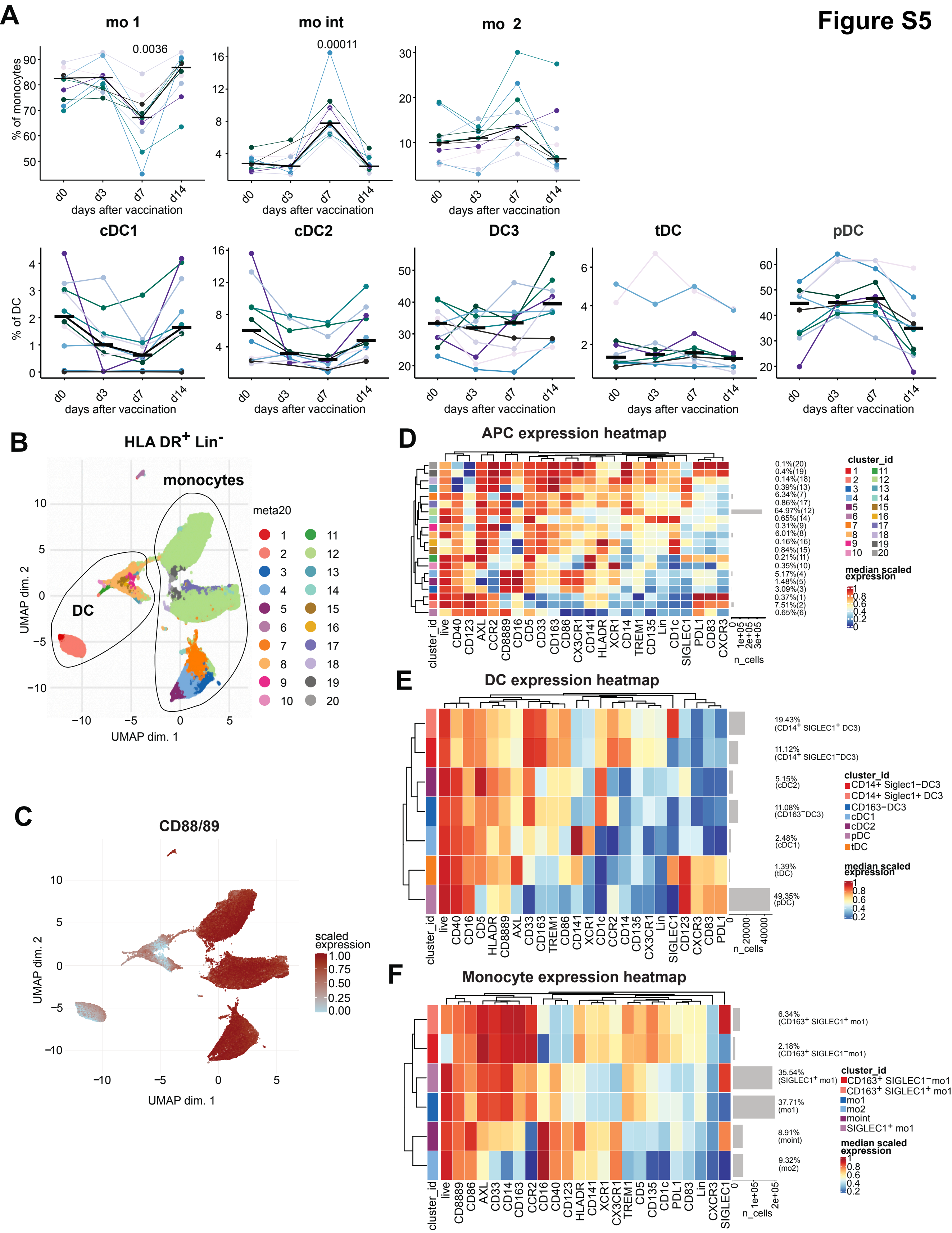
