## Supplementary Figure Legends for "Interferon-induced activation state of circulating dendritic cells and monocytes triggered by yellow fever vaccination correlates with early protective antibody responses"

**Supplementary Information**

### Legends to supplementary figures

#### Supplementary Figure S1

(A) Gating strategy for sorting APC subpopulations from the PBMC of vaccinees. B cells were gated as HLA-DR$^{+}$ Lin$^{+}$ CD19$^{+}$ and directly sorted (sort 1, vial 2). Monocytes were sorted as HLA-DR$^{+}$ Lin$^{-}$ CD88/89$^{+}$ (sort 1, vial 1) and then resorted into CD14$^{+}$ CD16$^{-}$ mo1, CD14$^{+}$ CD16$^{+}$ mo int and CD14$^{-}$CD16$^{+}$ mo2. DCs were sorted as HLA-DR$^{+}$ Lin$^{-}$ CD88/89$^{-}$ and then regated for HLA-DR$^{high}$ and separated into CD33$^{+}$ cDCs (sort 1, vial 3) and CD123$^{+}$ cells (sort 1, vial 4). Then cDCs were subsequently resorted into CD141$^{+}$ cDC1, CD141$^{-}$CD1c$^{+}$CD5$^{+}$ cDC2 and CD1c$^{+}$CD5$^{-}$ DC3. CD123$^{+}$ cells were resorted into Axl$^{+}$ CD33$^{+}$ tDC or double negative pDC. (B) Experimental design of bulk RNA seq experiment. Cells were sorted into PBMCs (100,000 cells) and subpopulations (1,000 cells). RNA was isolated using Quiagen RNeasy kits and used for adapted SmartSeqv2. (C) Expression heatmap of DC subtype-specific genes. Gene signatures were generated from DEGs between all the individual subpopulations (APCs, B cells and PBMCs) regardless of timepoint after vaccination. 25 highly expressed genes were selected for each population and the VST transformed expression values were visualized in the heatmap.

#### Supplementary Figure S2

(A) Normalized counts of IFN genes detected in bulk RNA sequencing data set shown for all populations over time. (B) Enrichment plots with normalized enrichment score of hallmark and reactome scores in DC3.

**Supplementary Figure 3**

Annotation of Louvain clusters obtained from scRNA-seq data. (A) Violin plots showing gene expression of DC and monocyte population markers. Shown is the expression level of the indicated genes in each Louvain cluster, shown by the color and the cluster number on the x-axis. (B) Stacked bars show the frequencies of the individual Louvain clusters before (d0) and at d3 and d7 after vaccination with YF17D. (C) UMAP visualization with annotated cell clusters shown as colored overlay, separated by individual donors and timepoints. (D) The dot plot shows the expression of the indicated genes in each Louvain cluster. Color intensities of the dots indicate the average gene expression values and diameters indicate the percentages of cells expressing the genes within the cluster. DC and monocyte population marker genes shown here are derived from our bulk sequencing dataset as well as from previous publications (Villani, Satija et al. 2017, Leylek, Alcantara-Hernandez et al. 2019).

**Supplementary Figure 4**

(A) Comparison of cDC1 d0 vs. d3 and d0 vs. d7 as well tDCs d0 vs d3 and d0 vs. d7 and moint/2 d0 vs d3 and d0 vs. d7 shown as volcano plots. Log2 fold changes are indicated on the x-axes and -log10 p values are indicated on the y-axes. The most significantly up- and downregulated genes (by p-value, log2 fold-change > 0.3) are annotated in the graphs.

**Supplementary Figure 5**

(A) The frequencies of monocytes and DC subpopulations over time were determined based on manual gating. Frequencies of mo1, mo int, and mo2 are shown as percentages of total monocytes. Frequencies of cDC1, cDC2, tDCs and pDCs are shown as percentages of total DCs (n = 10). Black horizontal lines indicate the median. Each time point after vaccination was compared to the baseline value. Significant p values are shown as calculated in R using the Kruskal-Wallis test and Dunn's multiple testing. (B) HLA $DR^+$ $Lin^-$ cells were exported from Flow Jo and Clustering analysis was performed on pooled samples of each of 3 batches from the 220 vaccinees. UMAPs of reclustered DCs with FlowSOM clusters indicated by colors are shown. DC and monocytes are annotated according to marker expression as found in B. (C) CD88/89 scaled expression indicated by color overlayed on the UMAP embedding. (D) Heatmap showing marker expression of each individual cluster in HLA $DR^+$ $Lin^-$ APC fraction. (E) Heatmap showing marker expression of clustered and annotated DC populations and of clustered and annotated monocyte populations.
